# Phylodynamic modelling of bacterial outbreaks using nanopore sequencing

**DOI:** 10.1101/2021.04.30.442218

**Authors:** Eike Steinig, Sebastián Duchêne, Izzard Aglua, Andrew Greenhill, Rebecca Ford, Mition Yoannes, Jan Jaworski, Jimmy Drekore, Bohu Urakoko, Harry Poka, Clive Wurr, Eri Ebos, David Nangen, Moses Laman, Laurens Manning, Cadhla Firth, Simon Smith, William Pomat, Steven Tong, Lachlan Coin, Emma McBryde, Paul Horwood

**Author notes:** ES, PH, EM, LC, ST planned and conceived of the study. ES conducted sequencing, wrote the code and conducted bioinformatic analyes; ES, SH conducted modelling analyses. IA, AG, RF, MY, JJ, JD, BU, HP, CW, EE, DN, ML, LM, CF, SS, WP, PH collected, maintained and provided the outbreak strains for sequencing, and managed all work in Papua New Guinea and Far North Queensland; ES wrote the draft of the manuscript, all authors contributed to the final version. authors contributed equally to the study (Emma McBryde, Paul Horwood).

## Abstract

Nanopore sequencing and phylodynamic modelling have been used to reconstruct the transmission dynamics of viral epidemics, but their application to bacterial pathogens has remained challenging. Here, we implement Random Forest models for single nucleotide polymorphism (SNP) polishing to estimate divergence and effective reproduction numbers (R_e_) of two community-associated, methicillin-resistant *Staphylococcus aureus* (MRSA) outbreaks in remote Far North Queensland and Papua New Guinea (n = 159). Successive bar-coded panels of *S. aureus* isolates (2 × 12 per MinION) sequenced at low-coverage (> 5x - 10x) provided sufficient data to accurately infer assembly genotypes with high recall when compared with Illumina references. *De novo* SNP calling with `Clair` was followed by SNP polishing using intra- and inter-species models trained on `Snippy` reference calls. Models achieved sufficient resolution on ST93 outbreak sequence types (> 70 - 90% accuracy and precision) for phylodynamic modelling from lineage-wide hybrid alignments and birth-death skyline models in `BEAST2`. Our method reproduced phylogenetic topology, geographical source of the outbreaks, and indications of sustained transmission (R_e_ > 1). We provide Nextflow pipelines that implement SNP polisher training, evaluation, and outbreak alignments, enabling reconstruction of within-lineage transmission dynamics for infection control of bacterial disease outbreaks using nanopore sequencing.

---

Sequence data from infectious disease outbreaks has provided critical information for infection control and inference of pathogen transmission dynamics, for example during the West African Ebola virus epidemic (1) and the current SARS-CoV-2 pandemic (2). Maximum-likelihood (ML) and Bayesian phylodynamic methods are commonly used to date the emergence of lineages and outbreaks, and to estimate key epidemiological parameters, such as changes in the effective reproduction number over time (R_e_) and the most recent common ancestor (MRCA) of an outbreak (3**?**, 4). Oxford Nanopore Technology (ONT) sequencing has emerged as viable technology for real-time genomic epidemiology, and has been applied across large-scale SARS-CoV-2 surveillance efforts in the United Kingdom and Denmark amongst others (5–8). Moreover, nanopore sequencing devices can be operated in low and middle-income countries where local genomics infrastructure may be lacking or is difficult to access (9, 10), so that a timely outbreak response is not feasible (11). This is particularly relevant for continuous surveillance at bacterial evolutionary time-scales, where outbreak strains may circulate for years, and can persist in human and animal reservoirs or the environment outside their hosts. Viral pathogen genomes, such as Ebola virus or SARS-CoV-2, are often sequenced directly from patient samples using targeted PCR-based enrichment approaches, achieving high genome coverage and resolution capable of informing phylodynamic models (1, 2). However, nanopore sequencing for bacterial pathogens, coupled to Bayesian phylodynamic models, have so far not been considered, mainly due the need for sufficiently accurate single nucleotide polymorphism (SNP) calling at bacterial whole genome scales (12). SNP calls from high coverage (> 30x) Illumina data is the current standard for accuracte SNP calls used in phylogenetic applications, but current generation nanopore SNP calling has suffered from low sequence read accuracy (R9.4.1) and a heavy focus on variant calling in human genomes, with much of the available callers developed specifically for human variants (13, 14). This problem is further aggrevated when attempting to sequence cost-effectively, e.g. using low-coverage multiplexed runs (< 5-10x) and simple library preparation with ONT sequencing kits (R9.4.1 pore architecture, SQK-RBK-004 libraries) that can be used in low and middle income countries with large burdens of bacterial disease.

Phylodynamic inference on nanopore platforms is further complicated when (ideally) using an outbreak reference genome that is closely related to the outbreak sequence type, thus providing sufficiently high variant calling resolution for transmission inference, particularly in recent transmisison chains or outbreaks (15). In addition, on bacterial time-scales (years) little sequence variation will have occured in newly emergent outbreaks, which places an disproportionate emphasis on correctly inferring the few available outbreak-specific polymorphisms. As a consequence, there is little room for systematic errors introduced by base- and variant callers when using (low-coverage) nanopore sequencing data to effectively survey bacterial outbreaks. Neural network-based, nanopore-native variant callers in particular can introduce excessive false positive SNP calls, complicating transmission inference from ONT sequence data, where accuracy and precision are required (16). Within-lineage phylodynamic inference for bacterial outbreaks additionally depends on sufficient temporal signal to ascertain a phylodynamic threshold, at which sufficient molecular evolutionary change has accumulated in the sample to obtain robust phylodynamic estimates (17–19). Due to slower substitution rates in bacteria compared to viruses (17), longitudinal sample collections are optimal for genomic epidemiology, and often require multiple years of data to infer transmission dynamics of the sampled population. Requirements for accurate whole genome SNP calls across a large number of isolates, sequenced cost-effectively at low genome coverage and over a sufficient interval of time, represents a significant barrier to the implementation of phylodynamic modelling for bacterial pathogens.

Illumina hybrid-corrected and ONT-native phylogenetic analyses methods have been demonstrated for a small number of distantly related bacterial nanopore genomes and genome assemblies from the same species e.g. *Neisseria gonorrhoeae* (16, 20) or between species from environmental sources (21). Recently a six-strain multiplex protocol for the MinION with genome assembly and determination of phylogenetic relationships to identify outbreaks has been tested for *S. aureus* lineages smapled in Norway. However, it remains unclear whether full within-lineage phylodynamic modelling is possible at population-level scale, whether estimates from nanopore data match results obtained using SNP calling with Illumina reads and whether sequencing runs can be conducted cost-effectively (at least 24 isolates per run). In this study, we adapt a variant polishing approach first implemented by Sanderson et al. (16) on metagenomic sequencing of *N. gonorrhoeae* using Random Forest classifiers to filter SNP calls from the nanopore-native variant callers `Medaka v1.2.3` and `Clair v2.1.1` (13). We use `Snippy` Illumina variant profiles as reference data and investigate caller performance across reference genomes and outbreak datasets. We show that Random Forest classifiers sufficiently remove incorrect calls from `Clair` in outbreak isolates with > 5x coverage to allow for sequencing of 24 community-associated *S. aureus* isolates per MinION flow cell (n_unique_ = 181) successfully resolving phylodynamic parameters of two outbreaks of ST93-MRSA-IV in remote Far North Queensland (FNQ) and Papua New Guinea (PNG).

## Results

We sequenced a total of 181 unique isolates from a paediatric osteomyelitis outbreak (collected between 2012 and 2018) in the Papua New Guinean highland towns Kundiawa (Simbu Province, n = 42) and Goroka (Eastern Highlands Province, n = 45). We additionally sequenced haphazardly collected blood cultures from a hospital in Madang (Madang Province, n = 8) and strains from routine community surveillance across Far North Queensland collected in 2019 (Cairns and Hinterlands, Cape York Peninsula, Torres Strait Islands, processed at Cairns Hospital, n = 86) (Fig. 1, Supplementary Tables). Oxford Nanopore Technoloy (ONT) sequencing was conducted using a minimal, dual-panel barcoding scheme, multiplexing 2 × 12 isolates interspersed with a nuclease flush on a single MinION flowcell (R9.4.1, EXP-WSH-003) for a total of 96 barcodes per outbreak (including isolate re-runs that were merged, n = 12, and external isolates excluded here, n = 3).

**Fig. 1.**
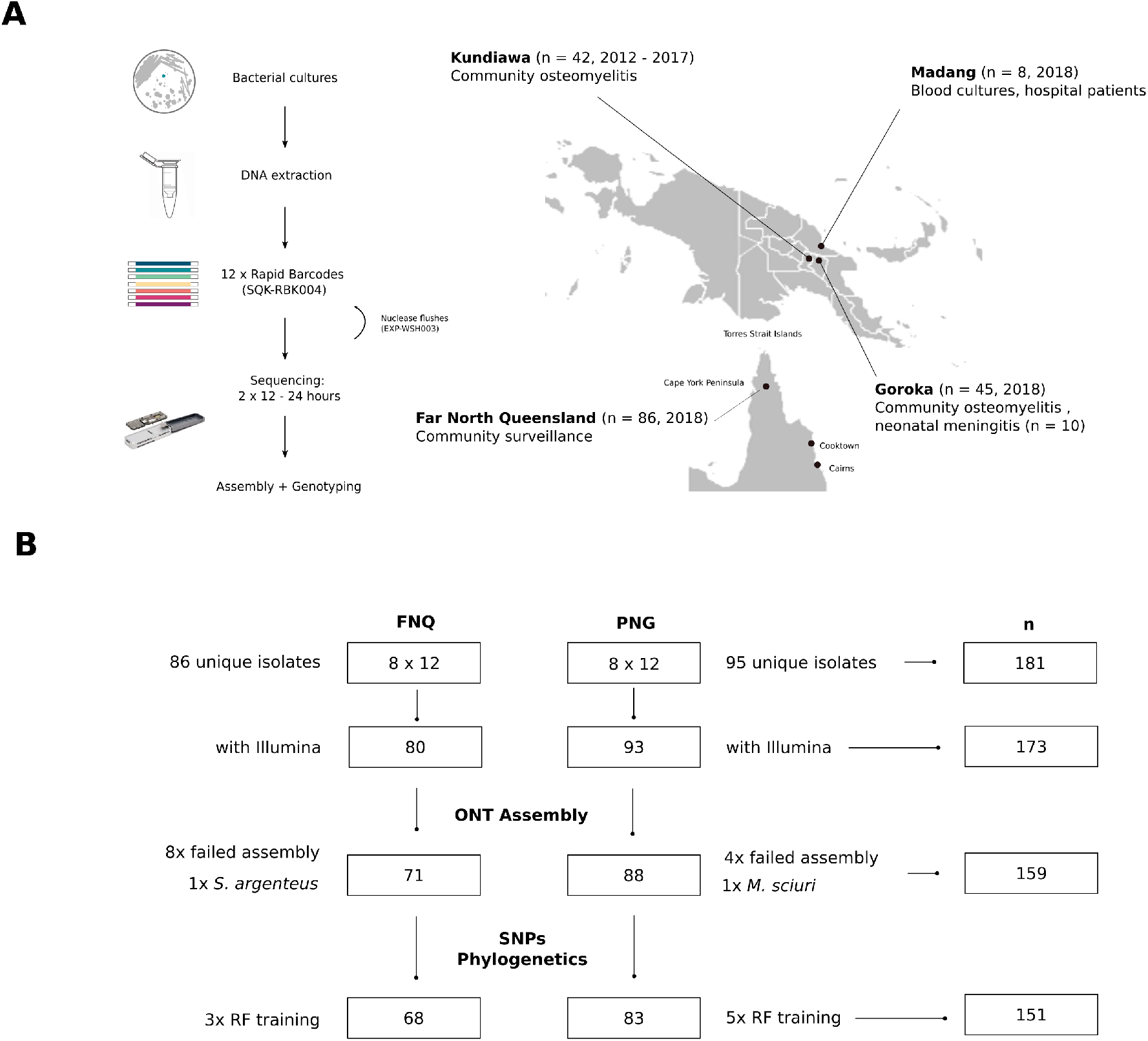
Sequencing workflow and outbreak sampling locations in northern Australia and Papua New Guinea (**A**). Isolates were sequenced on 8 flowcells with 24 isolates per flowcellu sing a sequential nuclease flush protocol. Sequenced data was subset to those matching Illumina sequencing of the isolates, assembled and quality controlled. Several isolates were set aside for independent Random Forest classifier training used in the SNP polishing and phylogenetics pipeline (**B**).

Rapid barcode sequencing libraries (RBK-004) were prepared omitting magnetic bead clean-ups after enzymatic digestion of cultured strains and simple spin column extraction. Panels produced between 0.506 - 6.47 Gigabases of sequence data per run (< 24 hours) resulting in low - medium coverage per isolate (ST93-JKD6159) (Fig. 2A). We excluded one infection with *S. argenteus* (FNQ) and one co-infection with *Mammaliococcus sciuri* (PNG). Isolates with matching Illumina data were retained to create a high-quality reference dataset for further evaluation of genome assembly and variant calling (n = 159, Fig. 1).

**Fig. 2.**
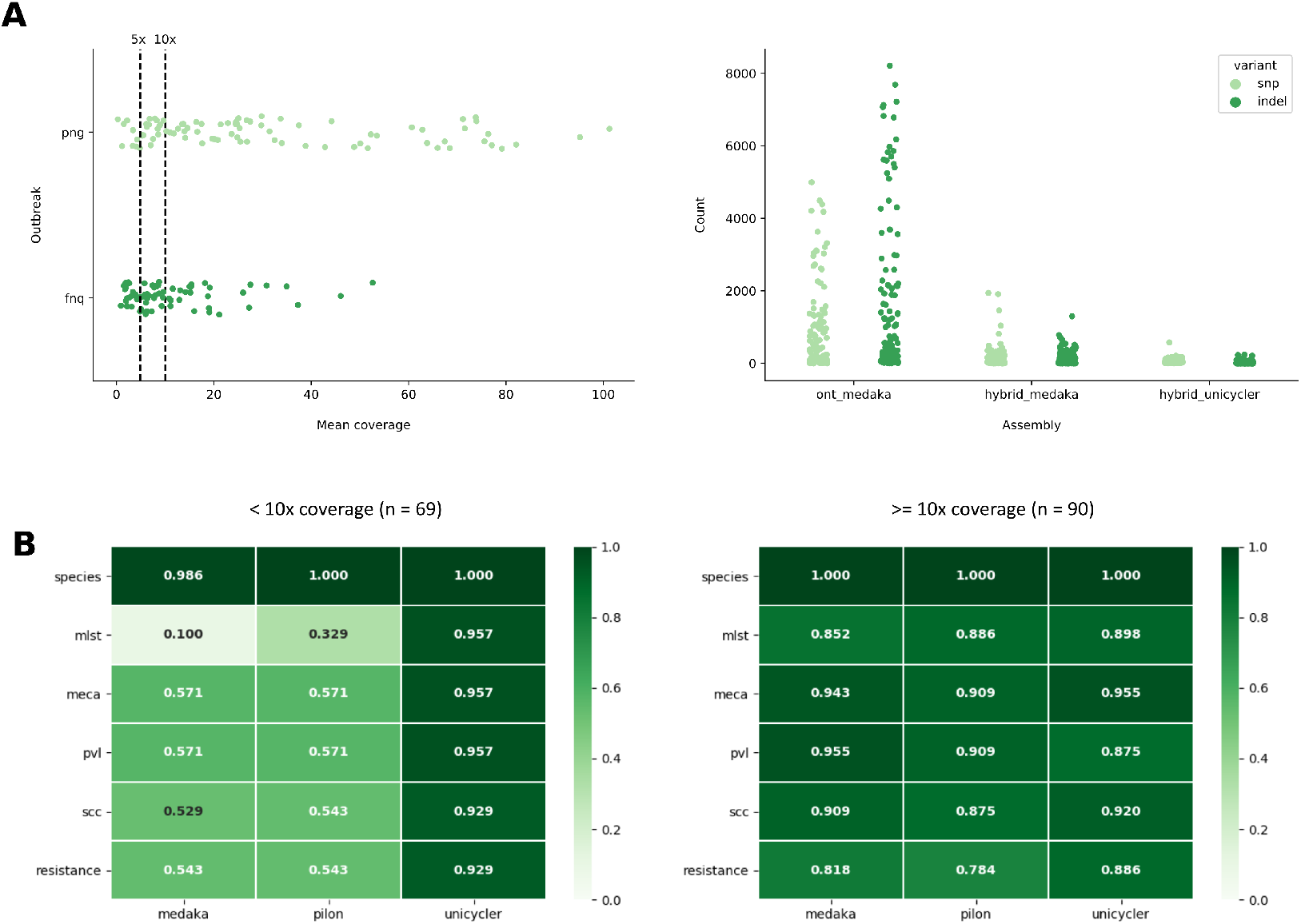
(**A**) Average genome coverage (R9.4.1, RBK-004) of `Bonito v0.3.6` base called nanopore reads against the JKD6159 (ST93) reference genome (n = 159) where the dashed lines indicate the coverage thresholds chosen to evaluate genotyping (10x) and phylodynamic models (5x) in the Far North Queensland (FNQ) and Papua New Guinea (PNG) outbreaks. SNP (light green) and indel (dark green) counts across three different assembly types: uncorrected nanopore reads polished with `Medaka` (ont_medaka), `Medaka` polished nanopore genomes Illumina corrected with `Pilon` (hybrid_medaka) and hybrid assembly in `Unicycler` (hybrid_unicycler) (**B**). Assembly genotyping results are shown as proportion of assemblies matching the reference Illumina genotype across the three types of assemblies, and the 10x coverage threshold.

### Genome assembly and genotyping validation

Short-read reference genomes, long-read polished nanopore genomes, and long-read hybrid genomes (`Pilon` (22) corrected long-read assemblies, `Unicyler` (23) short read assemblies with long-read correction) were assembled using a standardized `Nextflow` (24) pipeline wrapping `Shovill, Flye` (25) and `Medaka` and other components (Methods). Several isolates (12/159) failed long-read assembly due to excessive fragmentation of libraries and/or barcode attachment, but did not fail the short-read assemblies with `Skesa` (26) or the hybrid assemblies with `Unicyler` (Supplementary Tables 1-5), which first assembles short-reads and then scaffolds the assemblies with long reads to generate contiguous whole genome assemblies.

Compared to Illumina reference assemblies, SNP and indels were frequently occuring in low-coverage uncorrected nanopore assemblies (Fig. 2A, right). Errors were considerably reduced in high-coverage isolates leading to assembly identities ranging between 0.9993 and 0.9999 in the `dnadiff` metric (27) (Supplementary Tables 3, 4). Recovery of complete chromosomes and *S. aureus* specific genotypes from uncorrected long-read assemblies was sufficient for high-coverage isolates in our collection (Fig. 2B, > 80 −90%). Assembly genotyping for clinically relevant features such as the presence of *mecA* or the Panton Valentine leukocidin (PVL), major subtypes of SCC*mec* elements, resistance genes and other markers of interest showed high concordance with reference assemblies (Fig. 2B). In contrast, low-coverage assemblies often failed to call genotypes - recovery was low for *mecA* and SCC*mec* types, as well as for PVL and other markers of interest (Fig. 2B, < 60%, Supplementary Tables 3, 4). Hybrid long-read correction with `Pilon` did not markedly improve genotype recovery in low-coverage isolate; however, recovery improved in the `Unicyler` hybrid assemblies (Fig. 2A, 2B). Lower SCC*mec* subtyping performance was likely due to remaining insertions or deletions from nanopore data impacting on the large cassette chromosomes (> 20kb). `Unicyler` produced more accurate hybrid assemblies than correction of long-read assemblies with `Pilon` alone, and performed slightly better in hybrid assemblies of low-coverage nanopore data (Fig. 2B). For genome assembly and genotyping, our dual-panel sequencing approach recovers nanopore genotypes in high-coverage isolates (> 10x) although some errors remain, particularly in sequence type calling and SCC*mec* subtyping.

### Training and evaluation of Random Forest SNP polishers

Next, we aimed to accurately reconstruct the PNG and FNQ outbreaks within the maximum-likelihood background phylogeny of ST93. Subsequent phylodynamic analysis is challenging because accurate reconstruction of branch lengths within the nanopore clades is required for reproduction of the Bayesian epidemiological parameters. We first tried a candidate-driven approach, using Illumina core SNP panels from the ST93 background population (`Snippy`, n = 444, 6616 SNPs) and `Megalodon` which accurately reconstructed the divergence of the PNG clusters from the Australian East-Coast (Fig. S1). However, within-outbreak branch lengths were not reconstructed, because novel variation had accumulated since the divergence from the Australian east coast population in the 1990s (Steinig et al. 2021, *in preparation*). We therefore decided to use a *de novo* variant calling approach comparing two native nanopore variant callers based on neural network architectures, by default trained on *Home sapiens* variant calls (`Clair v2.1.1`) or a mix of human and microbial data from *Escherichiae coli*, *Saccharomyces cerevisiae*, and *H. sapiens* (`v1.2.3`). While recall was high, raw basecaller performance was exceedingly low in both `Clair` and `Medaka` accuracy and precision, particularly in outbreak isolate calls against the outreak reference genome (< 20%, Fig. S2).

We next adopted SNP polishers using Random Forest classifiers originally developed by Sanderson and colleagues (16) to correct nanopore variants in *Neisseria gonorrhoeae* from metagenomic data (Fig. 3, Methods). Each classifier was trained on three isolates with matching Illumina data and composite sequence features (Fig. 3B-D); because there were no considerations of specific training sets used in the original *N. gonorrhoeae* classifier, we trained *S. aureus* classifiers on three combinations of isolates including a mixed set of three sequence types (ST93, ST88, ST15) (saureus_mixed) and two sets of outbreak sequence type isolates (ST93) from either FNQ (saureus_fnq) or PNG (saureus_png). In combination with the original *N. gonorrhoeae* clasifiers, the different training sets allowed us to evaluate whether SNP polishing was effective using models from a different species entirely (sanderson), from the same species but without outbreak related data (saureus_mixed) or from the same species, but with isolates from the same sequence type or outbreak (saureus_fnq, saureus_png). All models trained on composite sequence features (Fig. 3, Methods) demonstrated high area under the curve (AUC) scores (0.976 - 0.989, orange) while models trained on quality features alone showed suboptimal AUC performance (0.748 - 0.760, blue) (Fig. 3B-D).

**Fig. 3.**
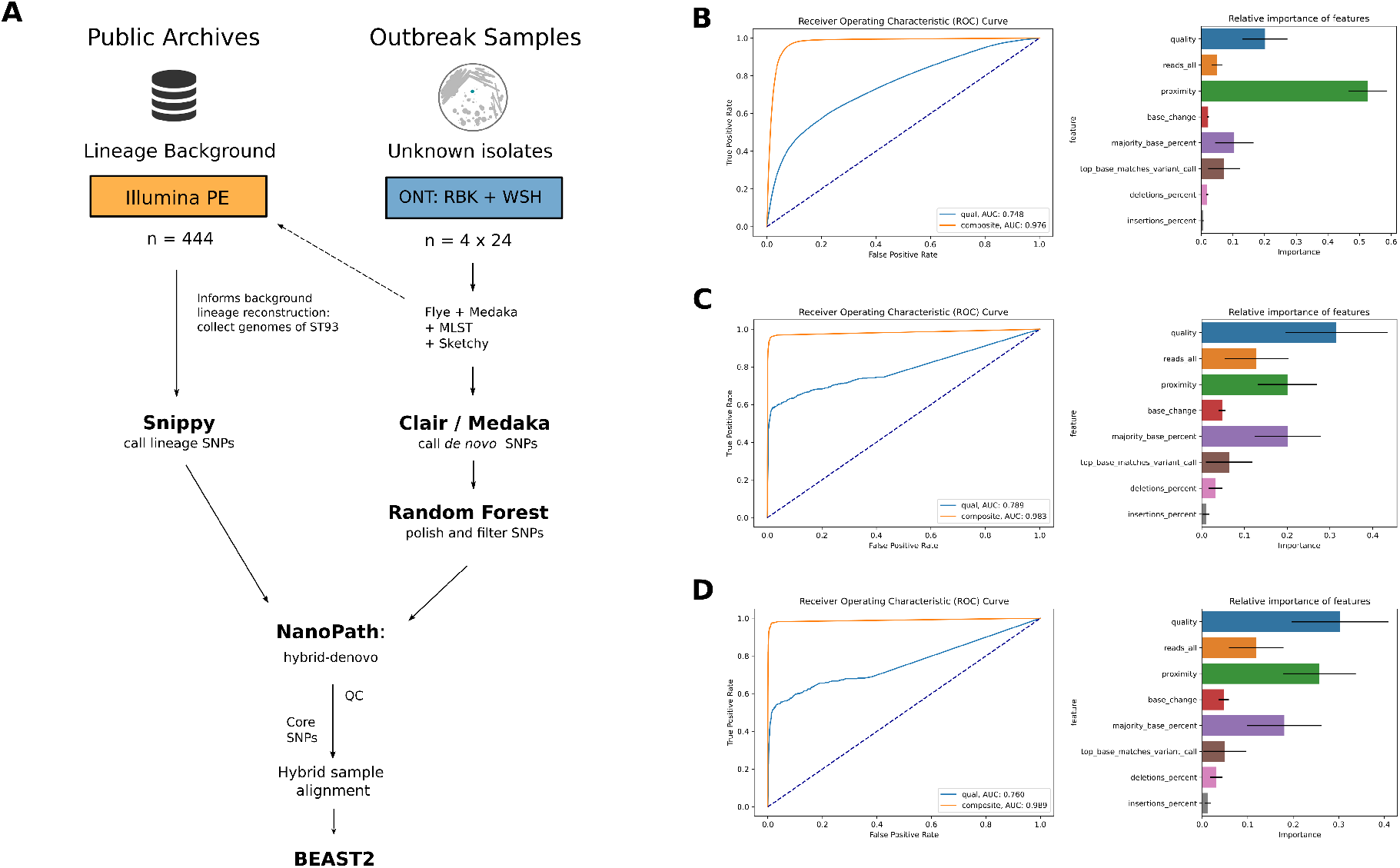
**A**) Workflow outlining culture-based protocol for community-associated *Staphylococcus aureus* nanopore sequencing using successive barcode panels on Oxford Nanopore Technology (ONT) MinION flow cells (R9.4.1). MLST typing informs the background population genome collection from a previous study (Illumina). Outbreaks in Papua New Guinea and Far North Queensland were caused by the Australian clone (ST93-MRSA-IV). SNPs are called for the Illumina background with `Snippy` and ONT outbreak isolates wth `Clair`. ONT SNP calls are polished using Random Forest SNP classifiers, trained on the outbreak reference genome (JKD6159 of ST93). (**B**) Area under the curve (AUC) scores of quality (blue) or composite (orange) features (left) used in training Random Forest classifiers for SNP polishing and relative feature importance of models (right) trained on (**B**) *S. aureus* mixed lineages (ST88, ST15 and ST93) (**C**) ST93 Far North Queensland isolates and (**D**) ST93 from Papua New Guinea with matching Illumina data and Snippy reference calls (all n = 3).

We next evaluated both the *N. gonorrhoeae* classifier, as well as the three *S. aureus* models against the remaining isolates from PNG and FNQ, excluding those used in training (Figs. 1B). Evaluations indicated that all trained SNP polishers increased accuracy and precision with slight reductions in recall (Fig. 4). However, sub-optimal performance was observed in all metrics for the *N. gonorrhoeae* classifier across outbreak sequence types (< 40%) as well as other sequence types (< 50%). Performance improved considerably in the mixed *S. aureus* polisher (saureus_mixed) both among outbreak isolates (69.52% ± 12.48*σ* accuracy, 75.94% ± 14.56*σ* precision) and other sequence types (81.94% ± 14.56*σ* accuracy, 90.11% ± 6.83*σ* precision). However, despite significant baseline improvement, the inter-species and mixed-sequence type models the number of false positive SNP calls remained in the range of 100s to 1000s (right column, Fig. 4A-B). Training the models with isolates from the same sequence type (ST93, FNQ) slightly improved performance (ST93: 71.69% ± 13.99*σ* accuracy, 83.33% ± 10.42*σ* precision) but reductions of accuracy and recall in other sequence types were observed (Fig. 4C). PNG outbreak-derived model (saureus_png) performed best for polishing isolates from the same outbreak across all metrics in the high coverage isolates (ST93: 69.28% ± 16.78*σ* accuracy, 87.57% ± 9.83*σ* precision) but incurred a steeper cost to accuracy and recall in non-outbreak isolates (Fig. 4D). Reductions indicate that the model trained on features specific to the outbreak genotype, and became significantly less generalizable to other sequence type applications. We note that the levels of precision and accuracy of the ST93 polishers in absolute numbers translate to 1 - 10s of false SNP calls compared to the *N. gonorrhoeae* and mixed seqeunce type model (Fig. 4).

**Fig. 4.**
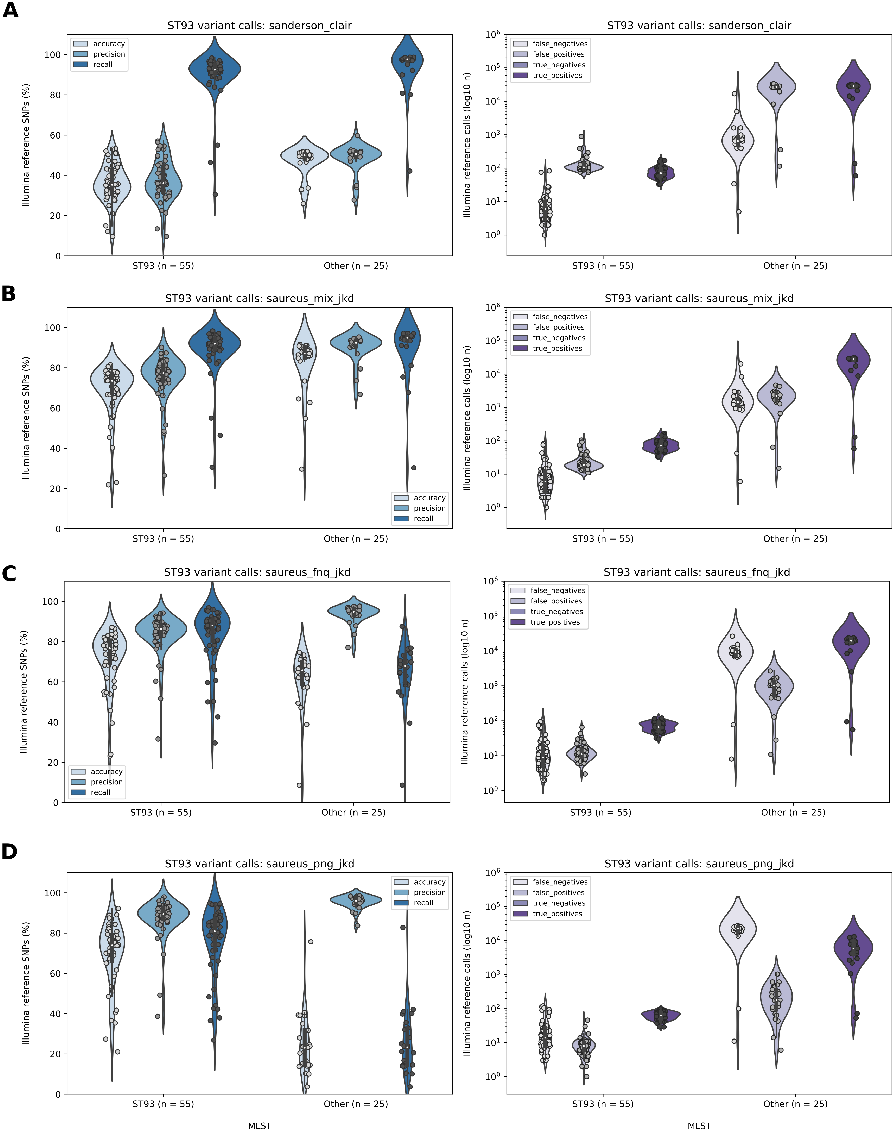
Trained Random Forest SNP polisher evaluation showing left: accuracy, precision and recall of `Clair` nanopore SNP calls against matching Illumina reference SNPs called with Snippy. Plots are split into ST93 outbreak isolates (inside left) and other sequence types (inside right) from Papua New Guinea (PNG) and Far North Queensland (FNQ) combined. In the right-hand plots the number of false negatives, false positives and true positive SNP calls for the groups is shown on a log-scale. Models were trained on three Illumina matched isolates from between-species (**A**) *Neisseria gonorrhea* from Sanderson et al. (16) within species (B) *Staphylococcus aureus* ST88, ST93, ST15 from PNG, (**C**) within-lineage (ST93) using samples from FNQ and separately from PNG (**D**, ST93). Polishing models were evaluated on all PNG and FNQ isolates excluding those used in training (ST93: n = 55, other sequence types: n = 25, > 10x coverage). Outliers in the tails of the distributions are novel multi-locus sequence type variants of ST93.

### Phylodynamic reconstruction using polished *de novo* SNPs

We next implemented `Snippy`’s core alignment functionality, calling sites present in all isolates of the sampled population, with a minimum SNP site coverage of 1x (JKD6159). Hybrid alignments integrated Illumina background SNPs from the ST93 (outbreak) lineage (n = 444) in combination with polished ONT nanopore calls from `Clair` (Fig. 1). The lineage background alignment, as one would use for short-read reference data, therefore served as a backbone for ONT data in the core-site alignment (Fig. 4B). We retained isolates with at least 5x coverage (n_5x_ = 531 / 562) due to low accuracy and precision of these isolates in the SNP polishing step (Fig. 4, Fig. S4). We then used the between-species, within-species, within-lineage (FNQ and PNG) models to apply for variant polishing in our *de novo* core alignment and phylodynamics pipeline (Fig. 3A).

`NanoPath`’s core alignment construction reproduced `Snippy`’s core alignment from Illumina data (6650 SNPs vs. 6662 SNPs, Fig.5A, B). When we called `Clair` SNPs on isolates with > 5x coverage from PNG (n = 56) and FNQ (n = 32) we observed a vast excess of SNP calls, particularly in the raw `Clair` calls, where the hybrid core alignment contained 491,210 SNP sites and was considered unusable (Supplementary Table 7). All polished SNPs produced reasonable alignments, where FNQ and PNG polishers produced alignments closest to the Illumina reference (Fig. 5, Supplementary Table 2). We reconstructed the ML phylogenies from these alignments in `1RAxML-NG` using the GTR+G model with Lewis’ ascertainment bias correction and rooted the trees on SRR115752 for comparison of topological consistency (28)). We also wanted to investigate whether the main introductions into FNQ and PNG could be reconstructed with accurate interpretations of their source divergence on the Australian east coast. For reference, we used Illumina alignments constructed with `NanoPath` (Methods) and `Snippy-core` with matching isolates (n = 531, Fig. 5).

**Fig. 5.**
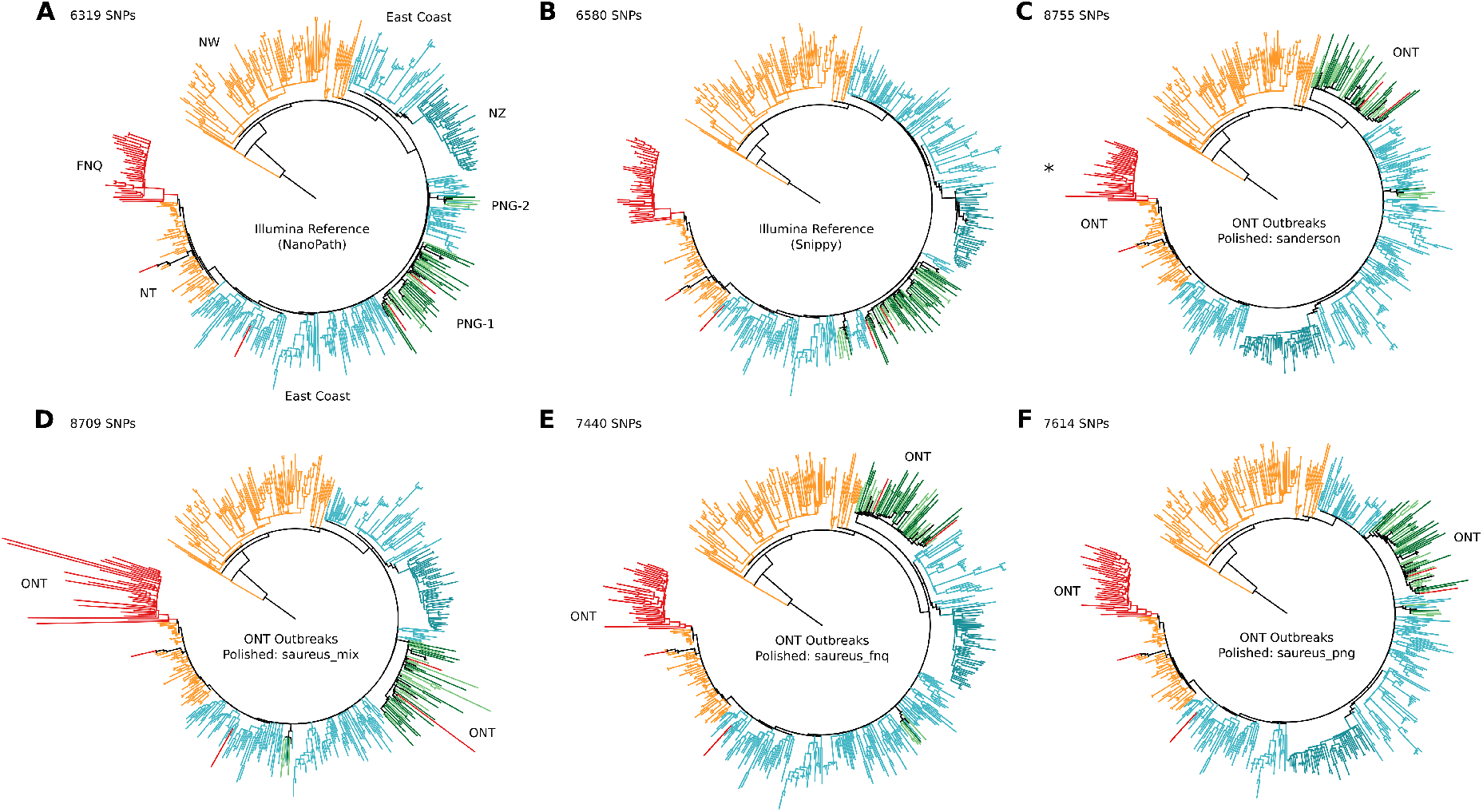
Comparison of maximum-likelihood phylogenetic topologies of ST93. Illumina reference trees were constructed with `NanoPath` (A) and `Snippy` (B). All other trees are hybrid phylogenies including the nanopore data of the outbreaks in Far North Queensland (FNQ) and Papua New Guinea (PNG) (> 5x coverage) within the ST93 background population (Illumina, n = 531) (**A)** after polishing `Clair` SNPs using the trained Random Forest classifiers (C: *Neisseria gonorrhoeae*; D-E: *S. aureus* mixed and lineage-specific). Asterisk (*) denotes two isolates with excessive branch lengths that were removed for visual clarity (Fig. S6).

All major clades and sub-populations of the background population (North West, East Coast, NT, and NZ) including the outbreaks in FNQ and PNG were accurately reconstructed as referenced by the Illumina trees (Fig. 5). Minor topological variations were observed in the position of the PNG-1 and PNG-2 introductions (greens), and the southern East Coast and NZ subclade (seagreen) of the East Coast population (turquoise, Fig. 5). However, there were no major topological inconsistencies that would affect interpretation of the source population. In all topologies, the outbreaks from PNG derived from the East Coast ST93-MRSA-IV clade, and the FNQ outbreak derived from the Northern Territory reintroduction (Fig. 5). Regional transmissions into the U.K. and Australia within the outbreak clusters remained identifiable (black and red branches in PNG-1 and PNG-2). Introductions into FNQ from other parts of the population are evident from both the reference and the polished alignments (red branches in East Coast, PNG and NT clades). Branch lengths of the nanopore-sequenced clades were similar to the reference ML tree, but were excessive in the between-species *N. gonorrhoeae* polished alignments as well as in the mixed sequence type alignments (Fig. 5, in particular due to two isolates: PNG-36 and PNG-62, Fig. S6). The alignment based on SNPs polished using outbreak seqeunce type (ST93) isolates were most consistent with the Illumina reference phylogeny of ST93. We note that within-lineage polishing did not require within-outbreak polishers, e.g. FNQ-trained polishers reproduced PNG outbreak divergence and vice versa.

We next investigated the performance of Bayesian phylodynamic methods to estimate the divergence date and effective reproduction number using birth-death skyline models with serial (PNG) or contemporaneous (FNQ) sampling and lineage-specific prior configurations (Steinig et al. 2020, *in preparation*). We ran `BEAST2` MCMC chains on the outbreak subsets of the full SNP alignment with sufficient isolates (n_PNG-1_ = 53; n_FNQ_ = 32) using a fixed substitution rate of the whole-lineage median posterior estimate (3.199 × 10^−04^). This was necessary as non-random sampling (subsetting the alignment to the outbreak clade) removes the temporal signal in the comparatively recent outbreaks, and thus leads to an overestimation of the outbreak tMRCA. We note that the models were efficiently run on a standard NVIDIA GTX1080-Ti GPU using `BEAST2` with the `BEAGLE v3` library at speeds of < 3-4 minutes / million steps in the MCMC (5-7 hours per run and GPU) making timely parameter estimation for outbreak responses feasible on low-cost hardware. On a NVIDIA P100 GPU, walltime decreased to < 50 seconds - 1 minute / million steps in the MCMC, around 1-2 hours walltime per run and GPU on a distributed system.

MCMC chains converged onto similar posterior distributions across all polished alignments in the PNG clade (Fig. 6). Polished models in the PNG clade were highly stable across posterior estimates, including those polished with between-species polisher from *N. gonorrhoeae*, and showing only slightly aberrant estimates of the MRCA in the mixed polishing model (Fig. 6B, Table S2). More variable posterior estimates were observed in the FNQ clade (Fig. 6), consistent with higher variability in branch lengths as a result of excessive false positive SNP calls retained in low-coverage FNQ isolates (Fig. 5). Neverthless, when compared to the `NanoPath` Illumina reference estimates, ST93-polished estimates (saureus_png, saureus_fnq) closely resembled those of the reference, with only minor deviations (Fig. 6, Table S2). Estimates were consistent with full lineage-wide analysis (R_e_ > 1.5-2.0, Steinig et al. 2021, *in preparation*) and we observed robust estimates in an exploration of the R_e_ prior (Table S2, Fig. S6, S7). We therefore demonstrate that SNP polishing enables the use of birth-death skyline models for outbreak parameter estimation, even with low-coverage nanopore sequencing data (5x - 10x). We have implemented training, evaluation and deployment of SNP polishers for within-lineage transmission modelling in `Nextflow`.

**Fig. 6.**
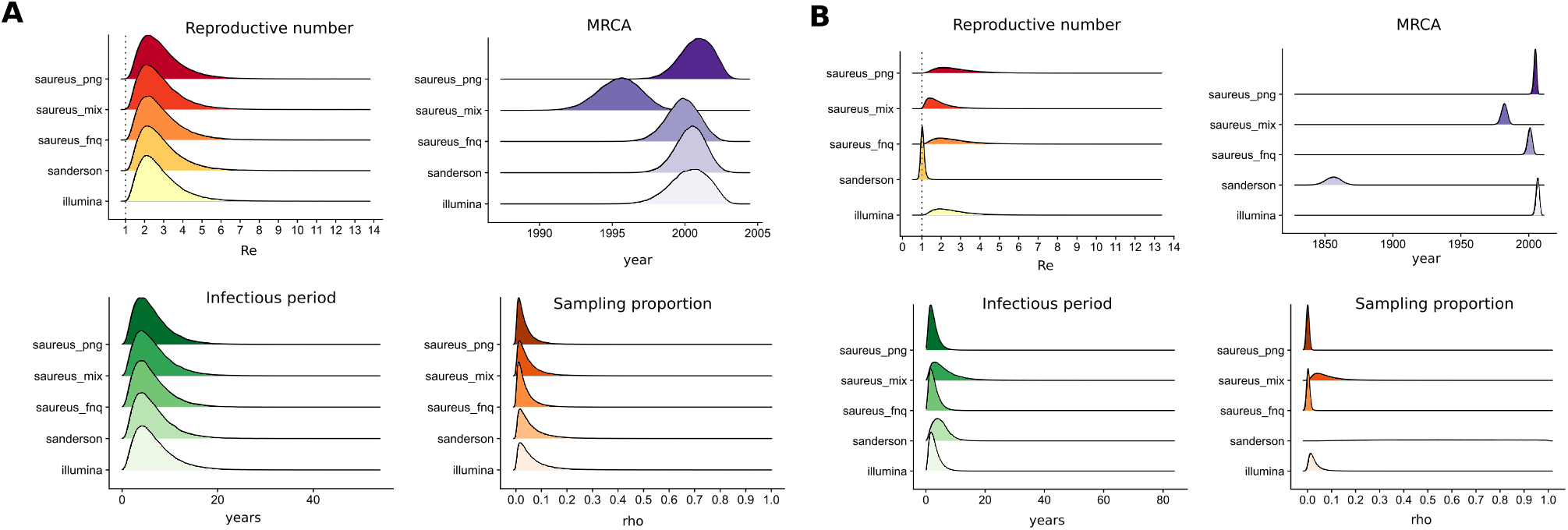
Posterior distributions of the effective reproduction number (R_e_, red), most recent common ancestor of the outbreak (MRCA, purple), infectious period (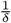, green) and sampling proportion (*ρ*, orange) for the nanopore sequenced outbreak clades in Papua New Guinea (**A**, PNG, n = 56) and Far North Queensland (**A**, FNQ, n = 32). Birth-death skyline models were run on the clade subsets of the polished hybrid alignments with > 5x coverage (ridge labels) including the Illumina reference alignment (illumina, bottom ridge), the between-species *Neisseria gonorrhoeae* polished alignment by Sanderson and colleagues (16) (sanderson), as well as the *Staphylococcus aureus* mixed lineages (saureus_mix), ST93 Far North Queensland (saureus_fnq) and ST93 Papua New Guinea (saureus_png).

## Discussion

In this study we provide a method for variant polishing and phylodynamic modelling of bacterial whole genome data using low-coverage nanopore sequencing. Previous studies using (high-coverage) nanopore data have evaluated phylogenetic reconstructions on few and distantly related isolates of *Neisseria gonorrhoeae* as well as other bacterial genomes from assembly (16, 20, 21). A recent pipeline for cluster identification using 6 strains per MinION flow-cell (42 on 7 flow-cells) successfully identified clusters in four distinct lineages, using a whole genome assembly based phylogeny (29). However, full outbreak reconstruction within the outbreak lineage — allowing for Bayesian model applications to estimate epidemiological parameters within the phylogeny — has so far not been conducted. Here, we show that the application of Random Forest SNP polishers developed by Sanderson and colleagues (16) can sufficiently reduce the number of false positive SNP calls from neural-network variant caller `Clair v.2.1.1` (13). Hybrid lineage alignments of ONT sequence and Illumina background data of the outbreak lineage (ST93) can be constructed, and effective reproduction numbers accurately modelled using birth-death skyline models in `BEAST2` (30). Interestingly, the Random Forest classifiers failed to polish `Medaka v1.2.3` reference-specific SNP calls (Fig. S3), even though the `Medaka-Bonito` model is trained explicitly on microbial signal data from *E. coli* and an experimental version (v0.1.0) was sucessfully used for polishing by Sanderson and colleagues (16). Polishing success of `Clair` calls suggest that the features selected here - in particular the proximity and quality features (Fig. 3B-D) - were effective at removing systematic false positive SNP calls, when trained with reference calls against specific reference genomes (e.g. ST93 outbreak genomes against ST93-MRSA-IV JKD6159 reference genome). Systematic error correction is supported by observations that SNP calling did not improve considerably using `Bonito v0.3.6` R9.4.1 DNA models compared to Guppy high accuracy (Fig. S5) and methylation-awar e models (data not shown). SNP polishers therefore appear to exploit systematic errors in the neural networks (trained on human variant calls) when applied to bacterial genomes. It remains to be seen whether re-training `Clair` or `Medaka` neural networks on *S. aureus* specific signal- and sequence-data would improve species-specific SNP calls.

We demonstrate the utility of our method by sequencing novel isolates of community-associated MRSA from a paediatric osteomyelitis outbreak in the highland towns of Kundiawa and Goroka (Papua New Guinea) and routine surveillance in remote northern Australia (Far North Queensland) (Fig. 1, n = 181). A protocol that minimised cost (without optimisation) allowed us to sequence two consecutive panels of 12 isolates with rapid barcoded libraries on a MinION flow cell (SQK-RBK-004), by using an interspersing nuclease flush (ONT, EXP-WSH-003). We note that spin column extractions resulted in several fragmented barcodes that failed assembly (12/96). Overall, phylodynamic models were mostly affected by very low coverage isolates (< 5x) whereas even low-medium coverage isolate (>= 5x) produced consistent estimates of the effective reproduction number for the PNG an FNQ clades, when compared to the Illumina reference (Fig. 6). Accurate modelling was possible even with inter-species polishers trained on *N. gonorrhoeae* in higher coverage isolates in PNG. Estimates were more variable in the low-coverage FNQ outbreak clade and for optimal performance some protocol optimisation will be required, and may include extraction protocols for long-reads, inclusion of a magnetic bead cleanup step (obligatory in the latest iteration of the ONT rapid kit protocols, May 2021), or short read elimination kits. While we were ultimately unable to use a total of 32 isolates (< 5x coverage) in the phylodynamic estimation, the cost per *S. aureus* genome using the 24x multiplex protocol ranges between USD $40 (no failures over 181 unique samples) and around USD $50 per genome with two repeat flow cells from already extracted cultures (Supplementary Material 2). Further optimization would incur small additional cost and can be conducted for bacterial pathogens of interest in sufficiently resourced laboratories. While we chose *S. aureus* as a model organism for this work mainly due to its relatively small genome (2.8 Mbp) and our interest in sequencing the outbreaks in PNG and FNQ, core principles and methods used in this study are immediately applicable to other bacterial pathogens and all steps of the pipelines are implemented in replicable `Nextflow` workflows (Methods).

We evaluated genotype reconstruction against previously sequenced Illumina data (Steinig et al. 2021) demonstrating the superior quality of hybrid assembly with `Unicyler`. Our genotyping analysis showed that for high coverage isolates (> 10x) genotyping directly from polished nanopore assembly was comparable to hybrid approaches (Fig. 2). We used the most recent models in `Bonito v0.3.6` for base calling followed by polished long-read assembly or hybrid assembly. With the imminent release of R10.3 pores and associated increases in raw read accuracy (estimated at Q20) we expect that the remaining misclassifications in genotypes from assemblies (mostly in MLST and SCC*mec* subtyping) will be eliminated and produce nanopore assemblies comparable to reference assemblies at > 5x - 10x coverage. We chose here to implement a rapid and minimal protocol to evaluate its application in remote reference laboratories, such as at the Papua New Guinea Institute of Medical Research. Our method requires some context from genomic surveillance at the level of full lineages (e.g. ST93 or ST772), in order to situate nanopore-sequenced outbreaks within the wider lineage context and fix the clade birth-death model substitution rate. Given that substitution rates vary between *S. aureus* lineages (17), an estimate from the background data is required to fix substitution rates within the outbreak clusters. For optimal polishing results it appears to be effective to train the Random Forest polishers on lineage-specific data, noting that effective polishing was still achieved when training isolates derived from a different part of the tree within the lineage (e.g. FNQ-trained polishers were effective on PNG isolates). In higher coverage isolates effective polishing was also achieved with the mixed *S. aureus* and *N. gonorrhoeae* models; we note that only three isolate with matching Illumina and ONT data are required for training the polishers.

We did not expect significant rate variation in the outbreak clades, which made computation of clade parameters with a lineage-wide fixed substitution tractable. We note that within-outbreak patterns of divergence vary between phylogenies (Fig. 5), and considering the number of remaining false positive and false negative SNPs after polishing (Fig. 4), we did not expect within-outbreak transmission chains to be reproducible. Optimization of SNP polishing or variant calling, for example with species-specific neural networks, remains to be investigated. For this study, we accelerated computation using the `BEAGLE v3` library (31) in combination with `BEAST2`. Moderate acceleration on standard hardware (< 5 - 7 hours) and increased acceleration on NVIDIA P100 GPUs (< 2 hours) were achieved. Nanopore-driven outbreak sequencing and GPU acceleration in `BEAST2` thus enable the rapid deployment of phylodynamic models and responsive surveillance of bacterial diseases.

## Materials and Methods

### Outbreak sampling in FNQ and PNG

We collected isolates from outbreaks in two remote populations in northern Australia and Papua New Guinea (Fig. 1). Isolates associated with paediatric osteomyelitis cases (mean age of 8 years) were collected from 2012 to 2017 (n = 42) from Kundiawa, Simbu Province (27), and from 2012 to 2018 (n = 35) from patients in the neighbouring Eastern Highlands province town of Goroka. We supplemented the data with MSSA isolates associated with severe hospital-associated infections and blood cultures in Madang (Madang Province) (n = 8) and Goroka (n = 12). Isolates from communities in Far North Queensland, including metropolitan Cairns, the Cape York Peninsula and the Torres Strait Islands (n = 91), were a contemporary sample from 2019. Isolates were recovered on LB agar from clinical specimens using routine microbiological techniques at Queensland Health and the Papua New Guinea Institute of Medical Research (PNGIMR). Isolates were transported on swabs from monocultures to the Australian Institute of Tropical Health and Medicine (AITHM Townsville) where they were cultured in 10 ml LB broth at 37°C overnight and stored at −80°C in glycosol and LB. Illumina short-read data from the ST93 lineage (28) included in this study were collected from the European Nucleotide Archive (Supplementary Tables).

### Nanopore sequencing and basecalling

2 ml of LB broth was spun down at 5,000 x g for 10 minutes and after removing the supernatant, 50 ul of 0.5 mg / ml lysostaphin were added to the tube and vortexed. Cell lysis was conducted at 37°C for 2 hours with gentle shaking followed by a *proteinase K* digestion for 30 mins. at 56°C. DNA was extracted using a simple column protocol from the DNeasy Blood & Tissue kit (QIAGEN) following the manufacturer’s instructions. DNA was eluted in 70 ul of nuclease-free water, quantified on Qubit, and DNA was stored at 4°C until library preparation. Library preparation was done using approx. 420 ng of DNA and the rapid barcoding kit with 12 barcodes (ONT, SQK-RBK004) as per manufacturer’s instructions. Basecalling was done using the R9.4.1 high accuracy (HAC, Fig. S5), the HAC methylation model (not shown), and the all context methylation odel (used for all analyses), run on a local NVIDIA GTX1080-Ti or a remote cluster of NVIDIA P100 GPUs. Sequence runs were conducted with 2 × 12 barcoded (SQK-RBK004) isolates per flowcell in two consecutive 18-24 hour runs. Libraries were nuclease flushed using the wash kit between consecutive runs (EXP-WSH-003). This is sufficiently effective to remove read carry-over, as demonstrated previously with hybrid assemblies of sequentially sequenced *Enterobacteriaceae* (32) and our analysis of a single library panel (FNQ-2) sequenced on a previously used flow cell with a human library. After washing with EXP-WSH-003 a total of 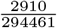 reads were classified as human in the *S. aureus* library, about twice as much as human contamination in other runs. Sequencing runs were managed on two MinIONs and monitored in `MinKNOW > v20.3.1`.

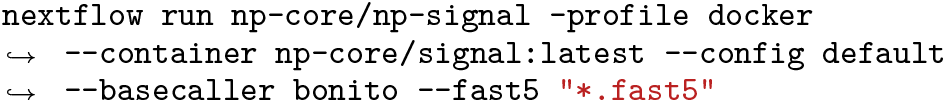

### Nanopore genome assembly and quality control

Genome assemblies for genotyping were constructed using our `Nextflow` assembly pipeline (https://github.com/np-core/np-assembly) which first randomly subsamples reads to a maximum of 200*x* coverage with `rasusa v0.3.0` (https://github.com/mbhall88/rasusa) and filtered *Q* > 7 with minimum read length of 100 bp using `nanoq v0.8.0` (https://github.com/esteinig/nanoq). `Fastp v0.20.1` (33) was used to trim adapter and low quality Illumina sequences. We then constructed three types of assemblies: a polished long-read assembly using ONT data only (flye), one with short-read correction of the ONT long-read assembly (pilon) and one that first assembles short-reads and scaffolds the assembly with long-reads. For the polished long-read assembly, `Flye v2.8.3` was used in conjunction with four iterations of `minimap2 v2.17-r941` (34) + `Racon 1.4.20` (35) and subsequent polishing with `Medaka 1.2.3`. For the long-read hybrid assembly, corrections were conducted with Illumina paired-end reads for each genome using two iterations of `Pilon v1.2.3`. For the short-read hybrid assembly, we used `Unicycler v0.4.8`. Reference Illumina assemblies were generated with the pipeline `Shovill v1.1.0` (https://github.com/tseemann/shovill) using `Skesa v2.4.0` and genotyped with `Mykrobe v0.9.0` (36) (from reads) and `SCCion v0.4.0` (https://github.com/esteinig/sccion), a wrapper around common assembly-based genotyping tools and databases (37–39) for *S. aureus*. We called species, resistance genes, virulence factors, Panton-Valentine leukocidin (PVL), multi-locus sequence type, *mecA* and major SCC*mec* cassette subtypes. We assessed differences between the Illumina references and hybrid- or nanopore assemblies using the `dnadiff v1.3` to determine assembly-based differences in SNPs and Indels, as well as assess overall identity between genomes (Supplementary Fig. 2). Coverage against the reference genome (ST93: JKD6159) (40) was assessed using `CoverM v0.6.0` (https://github.com/wwood/CoverM).

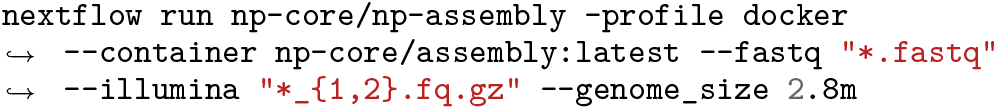

### *De novo* variant calling amd Random Forest SNP polishers

We called SNPs *de novo* using the neural-network callers `Medaka v1.2.3` (https://github.com/nanoporetech/medaka) and `Clair v2.1.1` (shown in example pipeline executions).

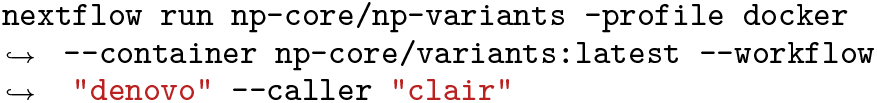

`Snippy v4.6.0` (https://github.com/tseemann/snippy) was used to generate a core site alignment of the ST93 background population (n = 444, 6161 SNPs) and reference Illumina core alignments including the outbreaks in FNQ and PNG isolates (> 5x, n = 531, 6580 SNPs). `Snippy` variant calls (SNP type) were used as reference truth for matching ONT and Illumina sequenced isolates.

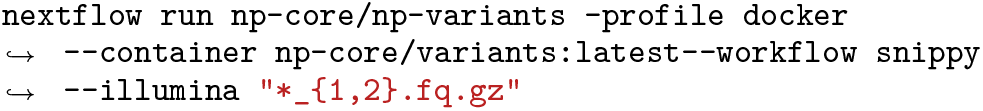

We implemented the feature extraction and Random Forest design from Sanderson and colleagues (16) who use the RandomForest classifier from `scikit-learn` (41) with default hyperparameter settings and feature extraction with `pysamstats`. Like the original implementation, we sub-sampled isolates to 2, 5, 10, 20, 50 and 100x coverage with `rasusa` to account for read coverage in training and evaluating the classifiers.. For training, we created three sets of matching Illumina and ONT sequence data, each with three isolates for training: three mixed sequence types (ST88, ST15, ST93) (saureus_mixed), one of Far North Queensland within-lineage isolates (ST93) (saureus_fnq) and one of Papua New Guinean within-lineage isolates (ST93) (saureus_png). Training and validation sets for the classifiers were split into 60% trianing and 40% validation data.

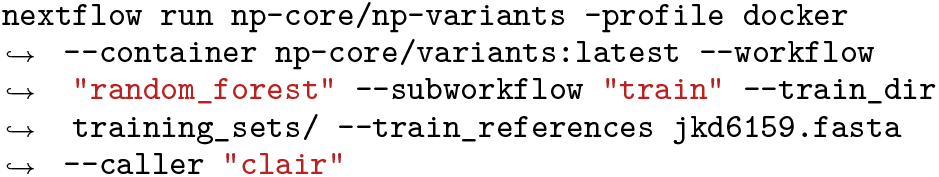

Next we evaluated the classifiers, including the *N. gonnorhoeae* classifier trained by Sanderson and colleagues, using the remaining isolates from FNQ and PNG (Fig. 1). We defined true positive (TP) SNPs as those that were called by both Illumina `Snippy` and ONT `Clair`, false positive (FP) as ONT SNPs that were not called with `Snippy`, and false negative (FN) `Snippy` calls that were missed by ONT calls or later excluded in the Random Forest filtering step. Since we used the *de novo* `Snippy` calls as reference, true negative (TN) calls (sites called as wild type by ONT and `Snippy`) were not able to be considered. We combined data from both outbreaks (n_ST93_ = 118, n_other_ = 44) and computed accuracy, precision, recall and F1 scores for each evaluation against Illumina reference data (Supplementary Tables 5, 6, Fig. 4).

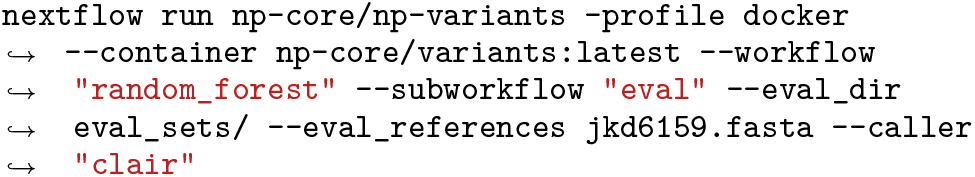

### Hybrid core site outbreak alignments

To contextualise polished ONT isolates called with `Clair` within the wider background of the ST93 lineage, we adopted the core functionality from `Snippy`’s core alignment caller (`snippy-core`) into an ONT and Illumina core SNP alignment caller in the `NanoPath` package (https://github.com/np-core/nanopath). Core SNP sites were defined by polymorphic SNP sites present in genomes of all isolates included in the alignment, excluding any site that in any one isolate falls into a gap, or any site with less than `--min_cov` coverage (default: 1x). We first polished ONT SNPs from `Clair` with the trained Random Forest models, including the *N. gonnorhoeae* dataset from Sanderson et al. (16). We then created reference alignments of the Illumina data (ST93 background and outbreaks, n = 531, > 5x) with `snippy-core`, as well as a reference Illumina and polished hybrid alignments with ONT outbreak SNPs in `NanoPath` (Fig. 5).

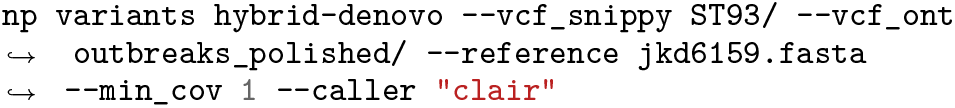

### ML phylogenetics and Bayesian model configurations

ML phylogeny of the ST93 lineage was reconstructecd from the Illumina and ONT polished alignments, including the outbreaks. We used `RAxML-NG` (42) with the general time reversible model and *Gamma* rate heterogeneity with 4 categories and the Lewi’s ascertainment bias correction for SNP alignments (GTR+G+ASC_LEWIS). Trees were rooted on SRR115236 (early isolate from 1992, near the root of the phylogeny) (28) and decorated with meta data of sample origin at state level in ITOL (43). Sampling dates in years were provided for each isolate.

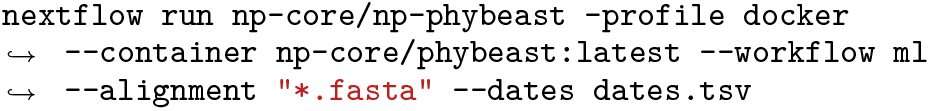

We next subsetted the full lineage alignments to the isolates in the large clades of the FNQ (n = 36) and PNG (n = 62) outbreaks. We then configured birth-death skyline models in `BEAST2` using a custom Python interface (`np beastling`) that stores model configurations of the serially (PNG) and contemporaneously sampled models (FNQ) in YAML files. Birth-death models consider dynamics of a population forward in time using the (transmission) rate *λ*, the death (become uninfectious) rate *δ*, the sampling probability *ρ*, and the time of the start of the population (outbreak; also called origin time) *T*. The effective reproduction number (R_e_), can be directly extracted from these parameters by dividing the birth rate by the death rate 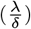. We configured the model priors as outlined in Table 1. Importantly, we set a lineage-wide fixed substitution rate prior (3.199 × 10^−04^, Steinig et al. 2021, *in preparation*) to account for the loss of temporal signal in the soutbreak subset alignments. `Beastling` constructs the `BEAST2` XML model files which can be run with the `BEAGLE` library on GPU:

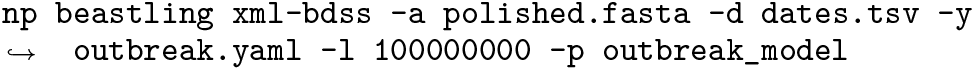

**Table 1.**
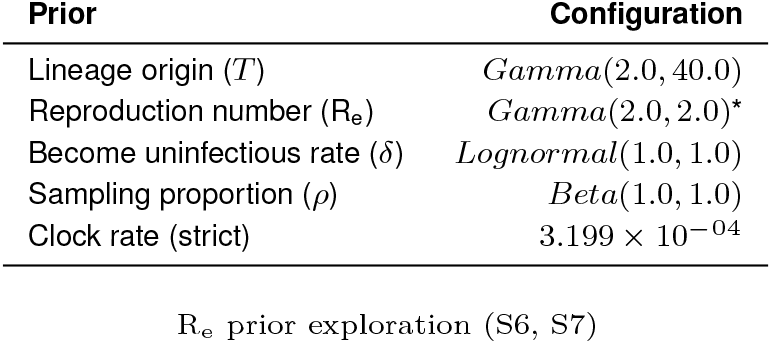
Birth-death skyline outbreak priors.

Results were summarized using the `bdskytools package` in R, where median higher posterior density intervals (HPD) were computed in custom plotting scripts that can be found along with all other results from the pipelines and model runs at: https://github.com/esteinig/ca-mrsa

## Data availability

Sequence data (Illumina, ONT) has been deposited under BioProject: PRJNA657380. `BEAST` XML files and logs of the model runs can be found at: https://github.com/esteinig/ca-mrsa

## ACKNOWLEDGMENTS

Es is supported by Queensland Genomics and a joint PRISM2 & HOT North (NHMRC ID: 1131932) pilot grant (44). CF is supported by a HOT North fellowship (NHMRC ID: 1131932). SYCT is supported by an Australian NHMRC fellowship (ID: 1145033). GPU models were run on the LIEF HPC-GPGPU Facility hosted at the University of Melbourne (LIEF Grant ID: LE170100200) (45).

## Supplementary Materials

### S1: Cost estimates of rapid dual panel barcoding protocol

- Sequencing cost per genome (as applied to data from PNG / FNQ, in Australian $): 4 × *FLOMIN* + 8 × *RBK* + 96 × *LS* + 96 × *DNeasy* + 96 × *Qubit* + 4 × *WSH* = $4, 882 at $50.22 per genome (excluding culture, tips etc.) and $53.29 per 181 unique genomes from PNG and FNQ
- Sequencing cost per genome (with resequencing of 48 isolates, in Australian $): 6 × *FLOMIN* + 12 × *RBK* + 96 × *LS* + 96 × *DNeasy* + 96 × *Qubit* + 6 × *WSH* = $6, 699 = approximate cost of $69.78 per genome (excluding culture, tips etc.) and $75.28 per 181 unique genomes from PNG and FNQ

### S2: Candidate-guided SNP calls using Megalodon

We evaluated a candidate-guided approach to reconstruct the phylogenetic divergences of nanopore-sequenced outbreak in PNG using a set SNPs at sites present in all isolates (core SNPs) called from existing population-wide background data of the ST93 lineage with `Snippy v4.6.0` (Illumina, n = 444, SNPs). SNPs from the known population were used as input to the candidate variant calling workflow in `Megalodon v2.2.10` (methylation-ware high-accuracy model, `Guppy v4.2.3`) and merged with the alignment of the background population (n = 495, SNPs). We used only isolates that passed genome assembly for the variant calling and phylogenetics (Fig. 1). Although slight variations in tree topology were observed in the divergence of the smaller monophyletic introduction into Papua New Guinea (PNG-2), the outbreaks diverged from their respective source populations (East Coast - PNG, North Eastern - FNQ) and deduction of their regional origin was not affected (Fig. S1). Importantly, putative transmissions from Papua New Guinea remained recognizable in the candidate-guided approach, thus allowing for the correct inference of sporadic regional transmission events (grey inside blue clade, Fig. S1). We estimated the date of divergence from the source population on the candidate phylogeny using `Treetime v0.8.1` (46) (PNG-1: 2004.45, 90% maximum posterior region (MPR): 2003.02 – 2005.71, PNG-2: 2000.74, 90% MPR: 2000.11 – 2001.47) and found that it reasonably approximated the estimate from the Illumina reference phylogeny (PNG-1: 2002.09, 90% MPR: 2000.97 – 2003.81, PNG-2: 2000.36, 90% MPR: 1999.73 – 2001.34). Estimates of lineage-wide substitution rates estimates from the SNP alignments were moderately consistent between the candidate approach (2.884e-04 +− 1.30e-05 *σ*) and the Illumina reference (3.174e-04 +− 1.19e-05 *σ*) and fall within the expected range of other *S. aureus* lineages and previous estimates for ST93. However, it was not possible to recreate within-outbreak relationships, because novel variation in the outbreaks was not captured in the core-genome variants of the background population (blue). Since the outbreak in the highlands has been ongoing since at least the 2000s, sufficient novel variation has accumulated in the PNG clade. Thus, given the absence of informative branch lengths within outbreaks, we were unable to conduct additional outbreak-specific phylodynamic analysis of these data.

**Table S1.** Reaction cost estimates (March 2021)

| Item | Reactions | \$AUD | per reaction |
| --- | --- | --- | --- |
| SQK-RBK-004 | 6 | 835 | 139.16 |
| FLOMIN-106D | 48 | 30,800 | 641.66 |
| EXP-WSH-003 | 6 | 110 | 18.30 |
| Lysostaphin | 315 | 835 | 3.15 |
| DNeasy | 250 | 1,728 | 6.91 |
| Qubit | 500 | 542 | 1.08 |

**Table S2.**
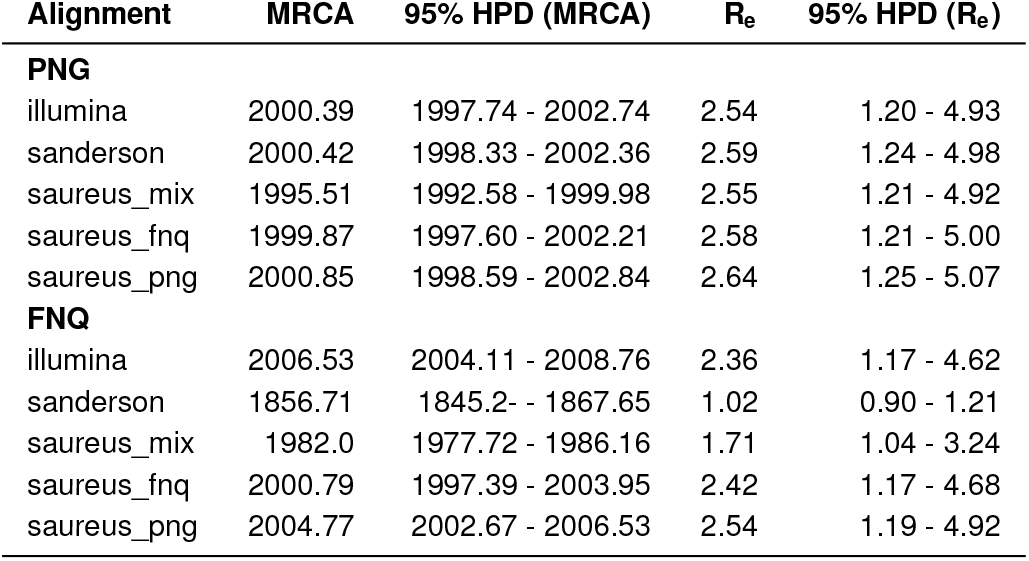
Birth-death skyline posteriors.

**Fig. S1.**
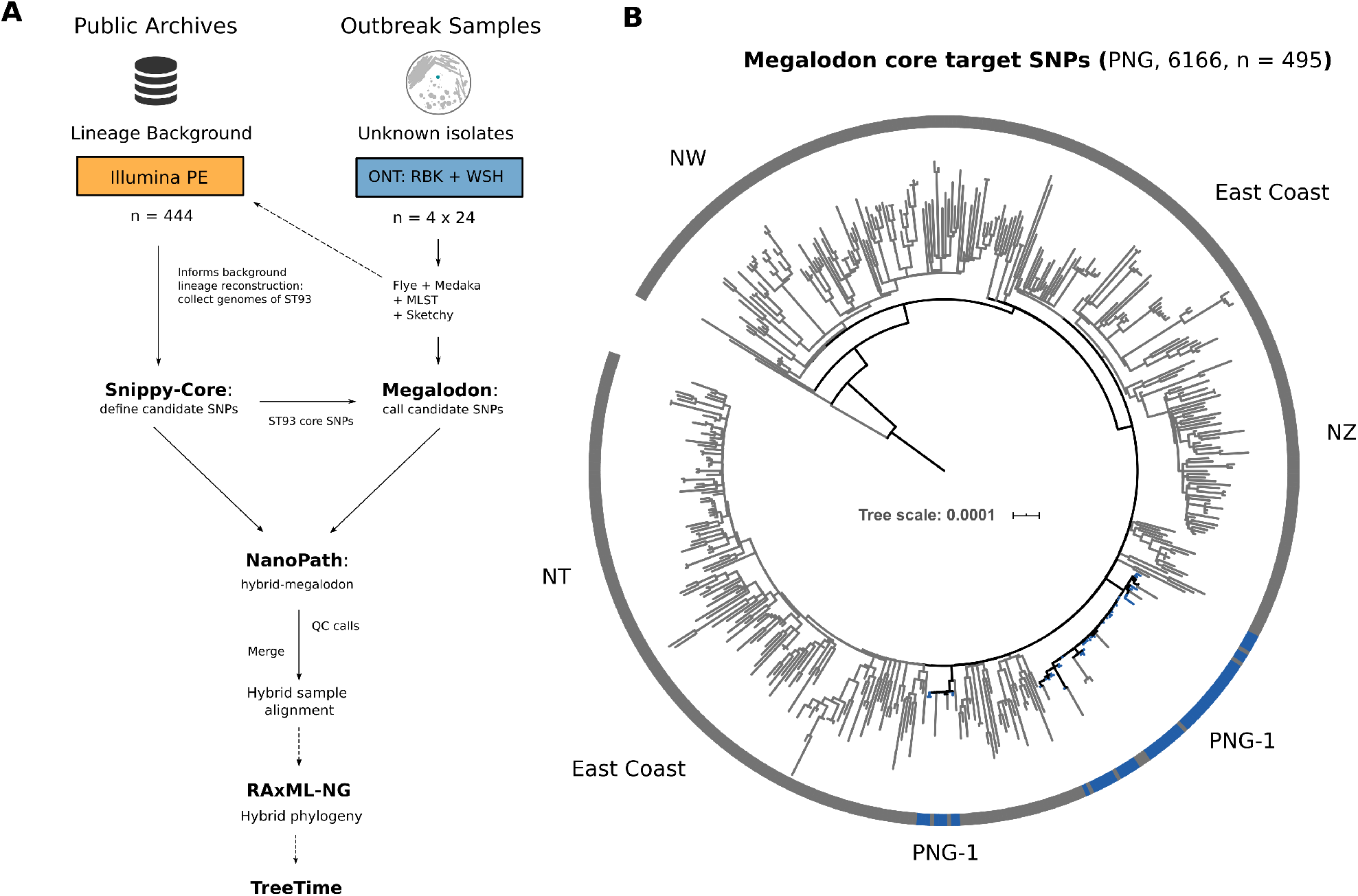
Candidate-guided variant calling workflow (A) and phylogenetic reconstruction of the Papua New Guinea (PNG) clusters PNG-1 and PNG-2 in the ML phylogeny (B) using candidate-guided core SNP sites from the lineage background population.

**Fig. S2.**
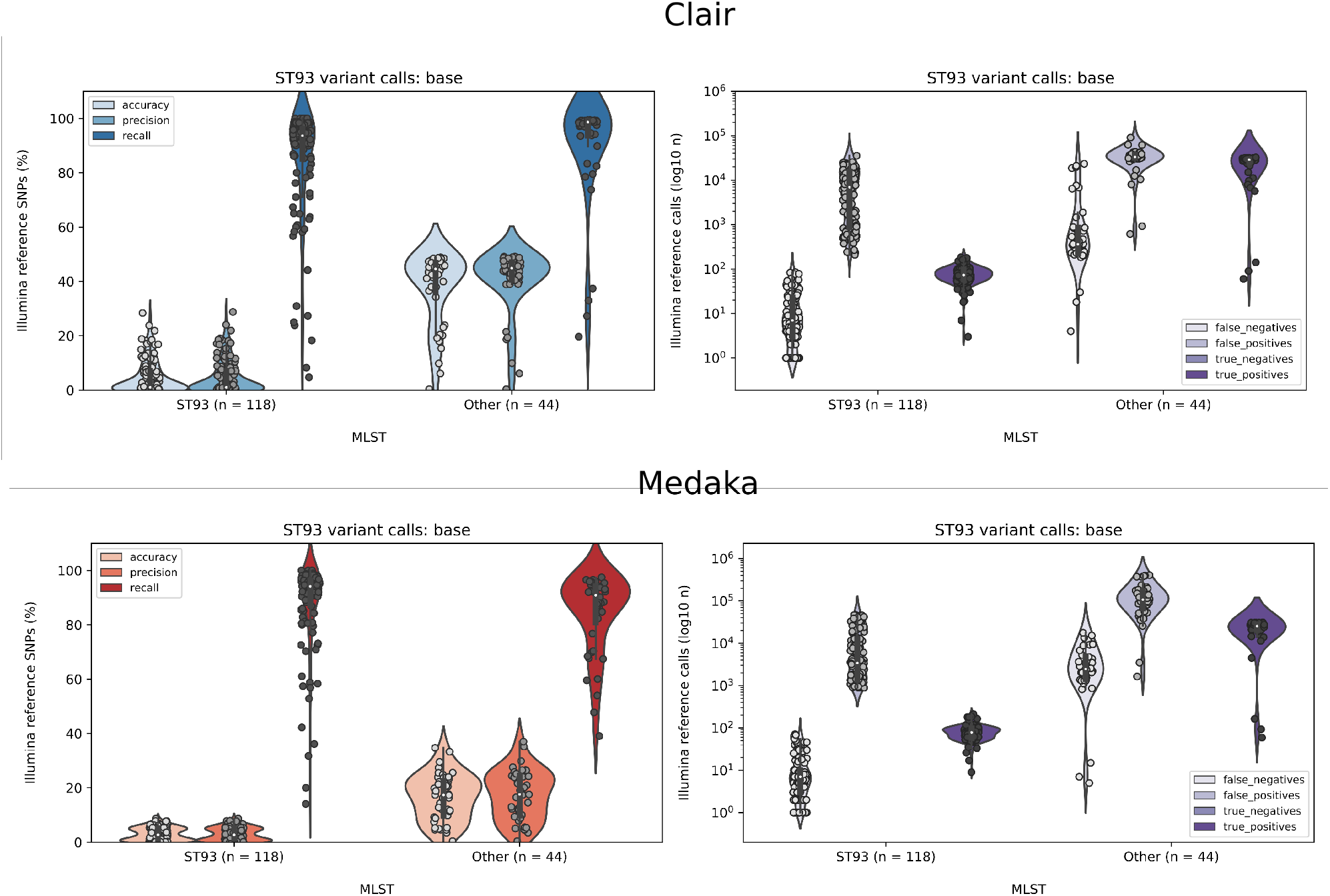
Raw SNP call accuracy, precision and recall (left, all isolates split into ST93 and other sequence types) from `Clair` (blue) and `Medaka` (red). Right plots show absolute numbers of false negatives, false positives and true positives on a log scale.

**Fig. S3.**
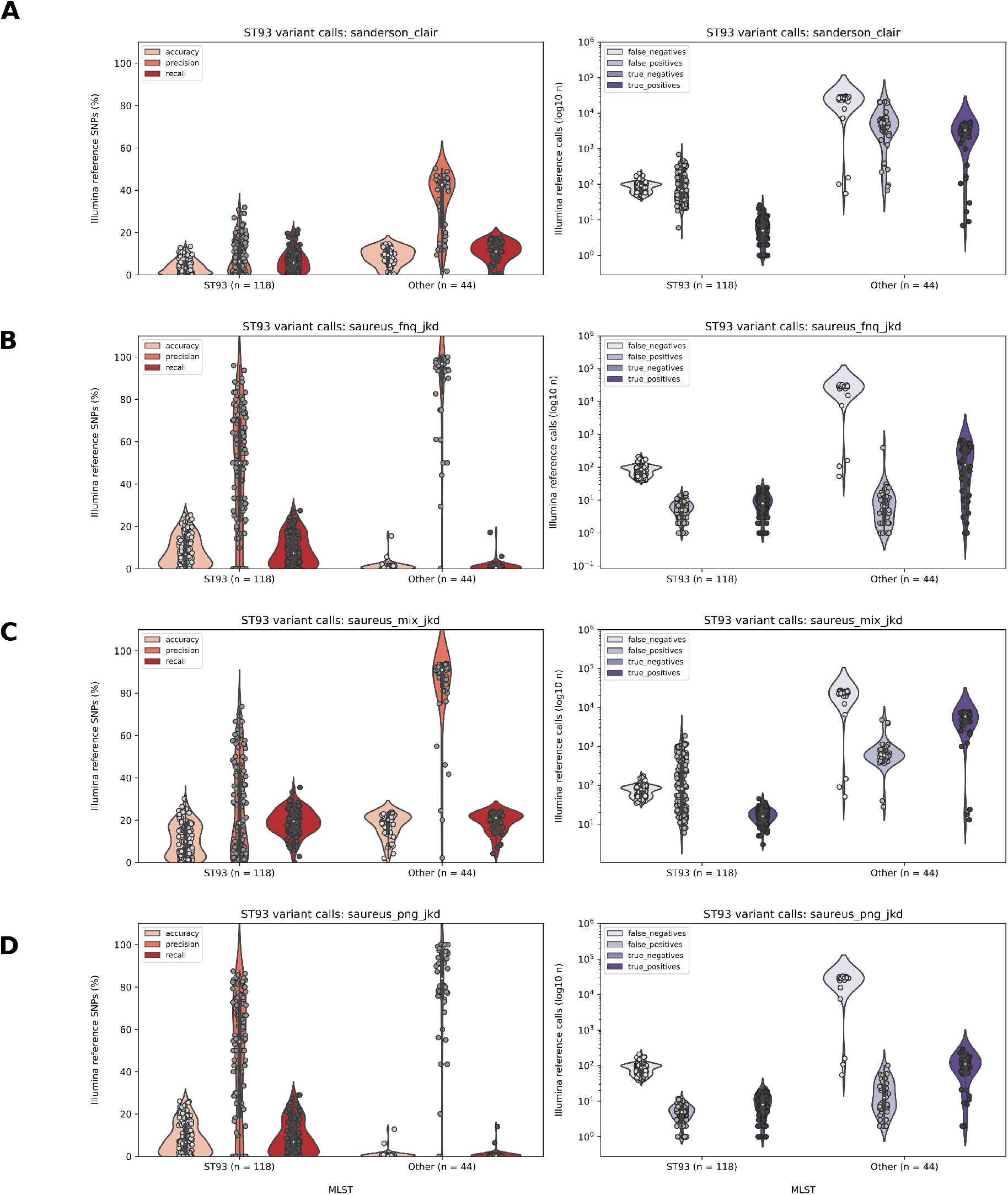
`Medaka` (all coverage) SNP calling accuracy, precision and recall compared to Snippy (Illumina reference) calls. Left plots show the metric distribution across ST93 (outbreak clades, n = 118) and other sequence types (split panels, n = 44). Right plots show absolute numbers of false negatives, false positives and true positives on a log scale.

**Fig. S4.**
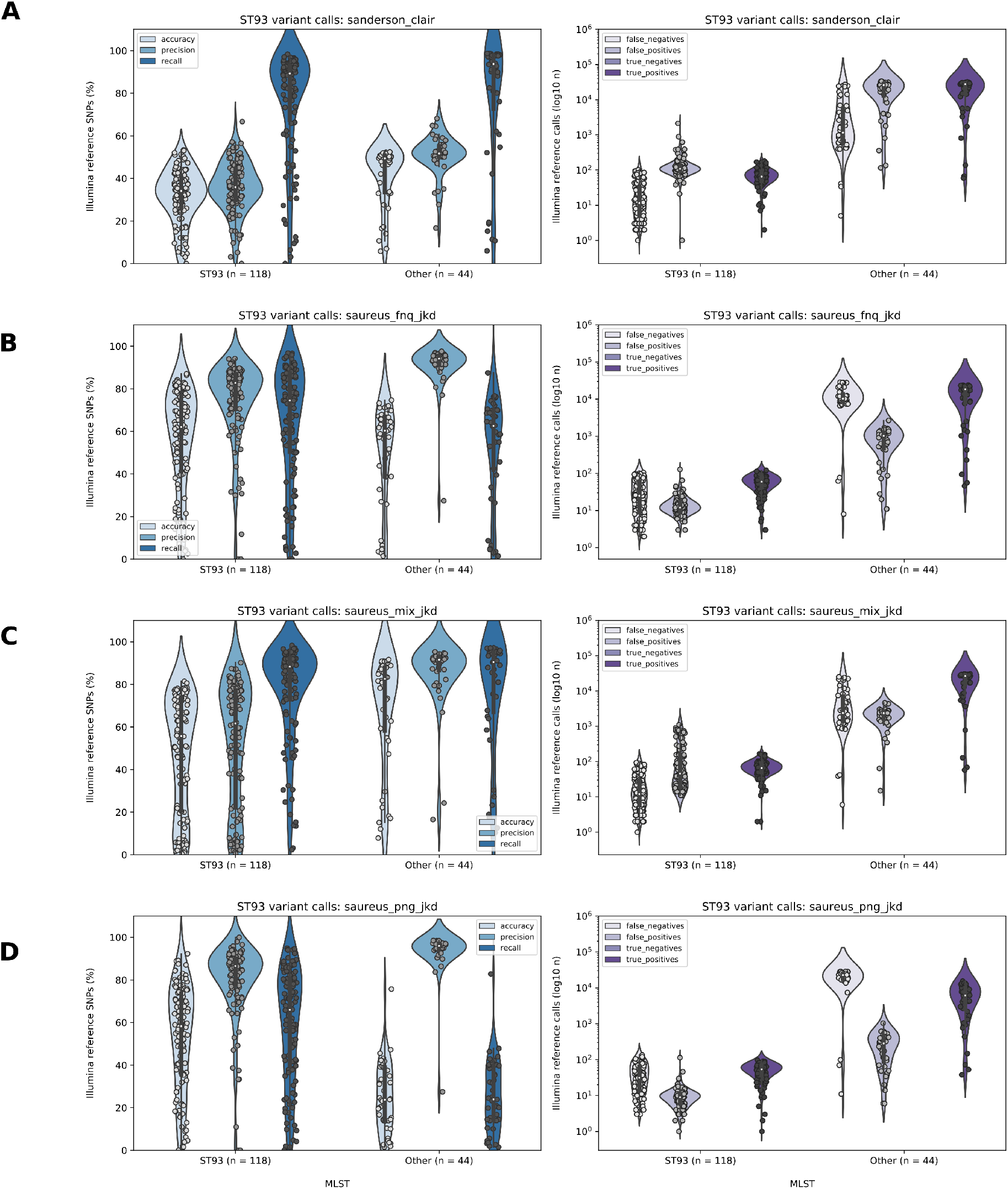
`Clair` (all coverage, n = 159) SNP calling accuracy, precision and recall compared to `Snippy` (Illumina reference) calls. Left plots show the metric distributions across ST93 (outbreak clades) and other sequence types (split panels). Right plots show absolute numbers of false negatives, false positives and true positives on a log scale.

**Fig. S5.**
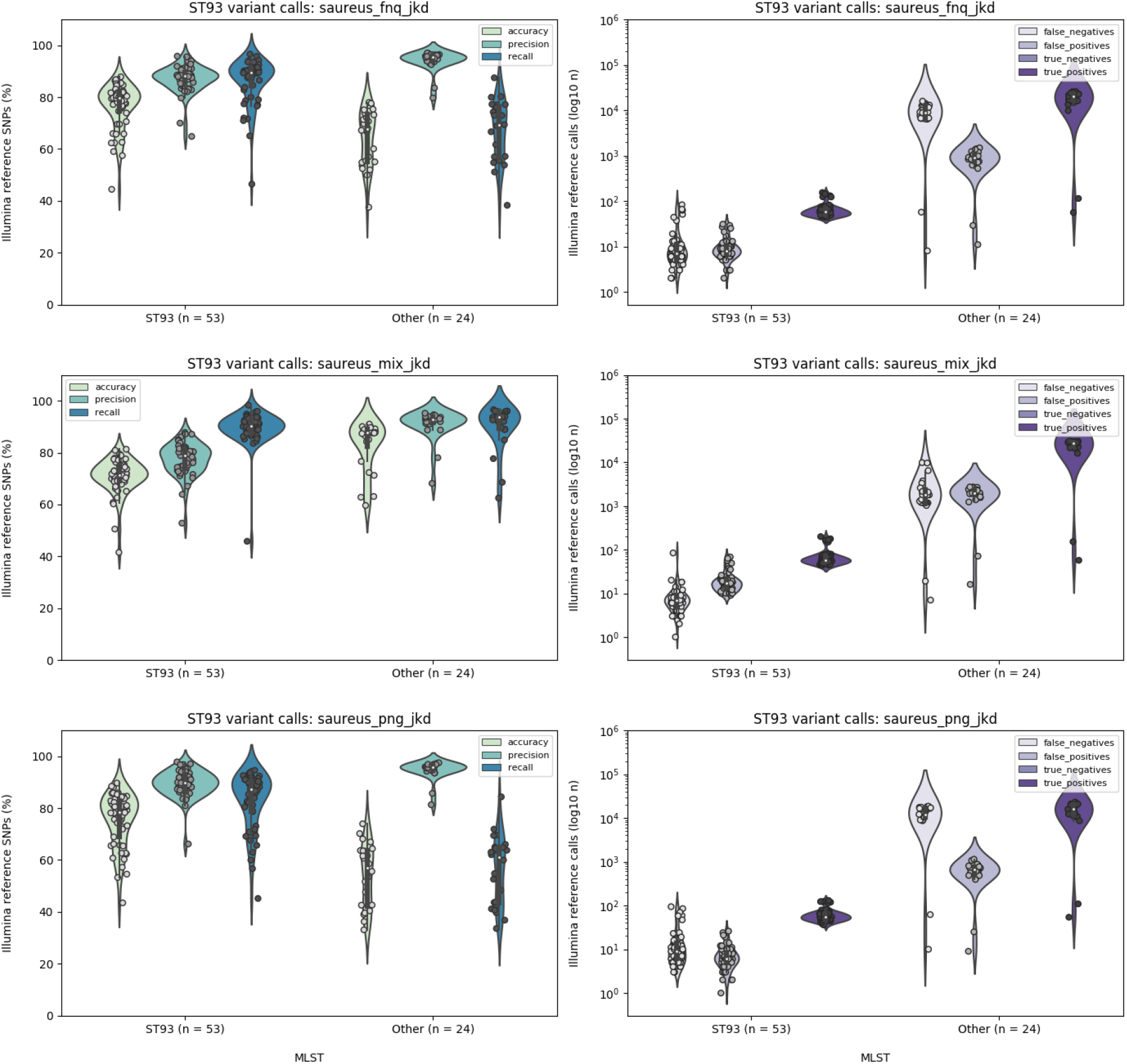
Isolates from the Papua New Guinean outbreak polished using Random Forest classifiers on `Guppy v.4.2.3` (high accuracy model) base called reads and SNP calls using `Clair`, showing similar error profiles as `Bonito v0.3.6` base called reads and SNP calls.

**Fig. S6.**
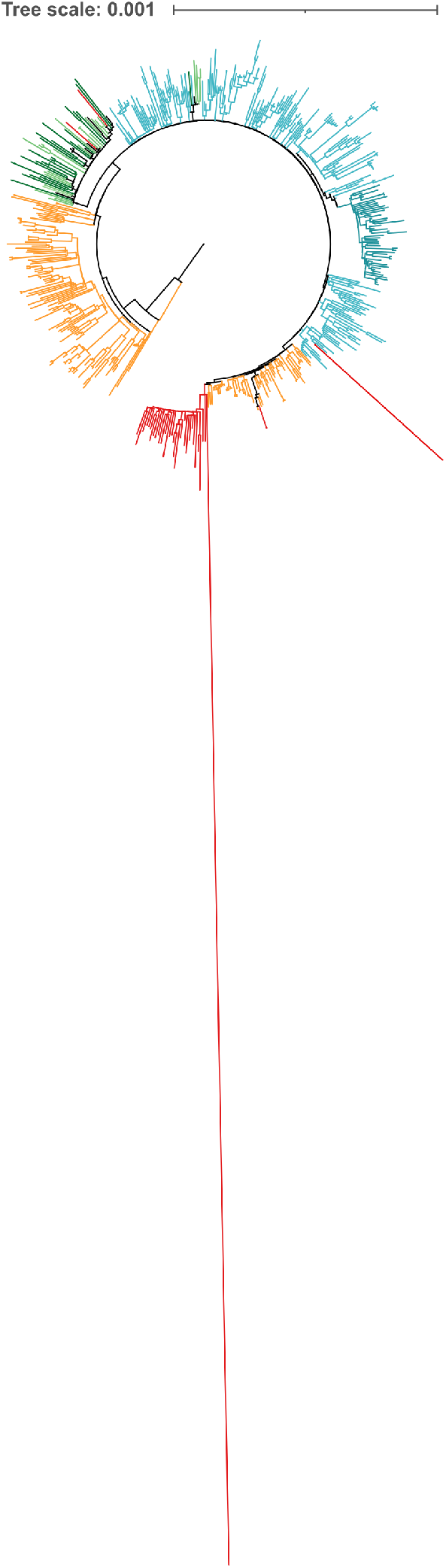
Maximum likelihood phylogeny of *Neisseria gonnorhoeaea* polished ONT SNPs of ST93-MRSA-IV outbreaks in Far North Queensland (FNQ, red) and Papua New Guinea (green). Complete branches are shown compared with Fig. 5, including extremely abnormal branch length of FNQ-36 and FNQ-62.

**Fig. S7.**
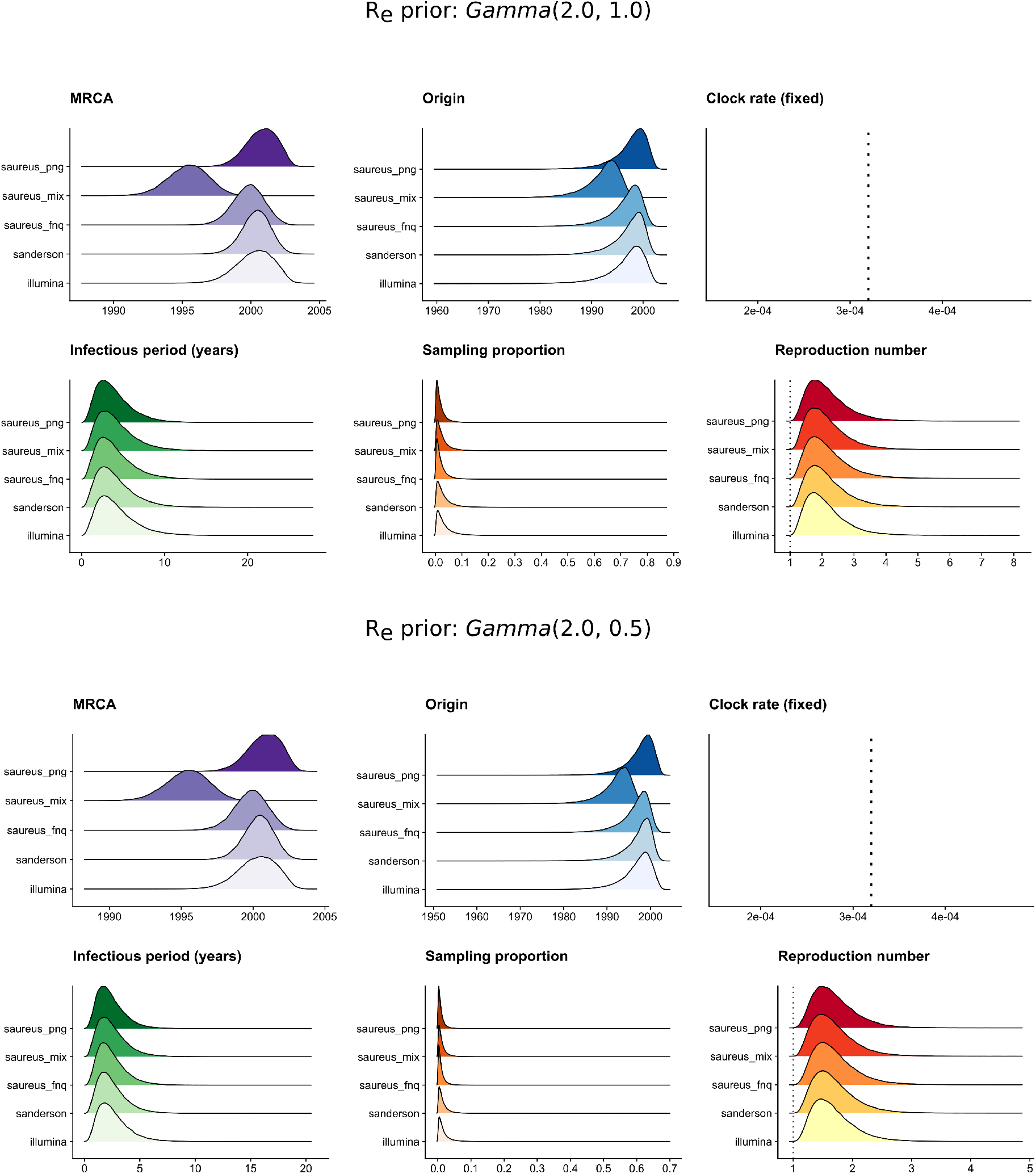
Birth-death skyline posterior estimates of the serially sampled PNG outbreak of ST93-MRSA-IV with a different prior of the effective reproduction number (R_e_) using a *Gamma*(2.0, 1.0) and *Gamma*(2.0, 0.5) configuration.

**Fig. S8.**
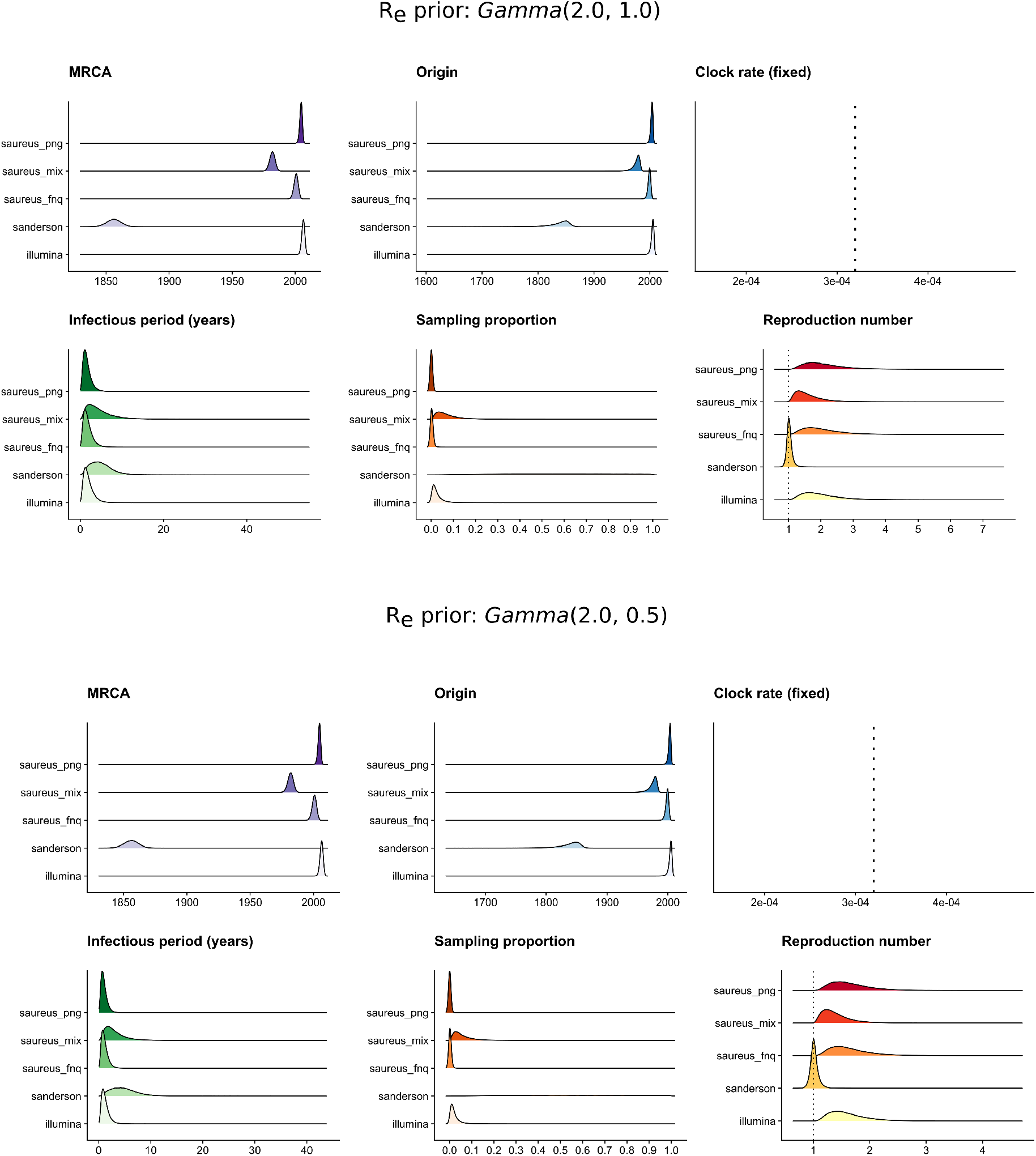
Birth-death skyline posterior estimates of the contemporaneously sampled FNQ outbreak of ST93-MRSA-IV with a different prior of the effective reproduction number (R_e_) using a *Gamma*(2.0, 1.0) and *Gamma*(2.0, 0.5) configuration.

